# NFATc2 potentiates DNA double-strand breaks repair by interaction with Ku80 in radiotherapy patients

**DOI:** 10.64898/2026.07.30.741465

**Authors:** Théo Barthélemy, Joshua Dulong, Lou Riedel, Sandra Moratille, Nicolas O. Fortunel, Jérôme Lamartine

## Abstract

A fraction of patients treated with radiotherapy are known to be more sensitive to ionizing radiations. Skin fibroblasts from such radiosensitive individuals exhibit a higher cellular toxicity after irradiation and a delay in DNA repair. Deciphering the molecular mechanisms underlying these cellular defects is thus of major importance. We previously observed that the transcription factor NFATc2 is expressed at a reduced level in fibroblasts from radiosensitive patients. The present work aimed to elucidate the role of NFATc2 in the regulation of DNA repair, particularly the repair of radiation-induced double-strand breaks. We demonstrate an interaction of NFATc2 with the NHEJ repair protein Ku80 and observe that the NFATc2 RHD domain is necessary and sufficient for this interaction. Moreover, we show that NFATc2-Ku80 complexes are not colocalized to DNA double-strand breaks sites suggesting an involvement upstream of the DNA repair pathway. The silencing of NFATc2 impairs the NHEJ repair activities by delaying Ku70-Ku80 interaction in the early steps of this pathway. Finally, stable over-expression of NFATc2 in patients’ fibroblasts partially rescues their defective DNA repair phenotype, especially in the most radiosensitive cells. Altogether, our data reveal that NFATc2 is a regulator of DNA repair in skin fibroblasts and therefore a potential modulator of cellular radiosensitivity.

## Introduction

Among DNA damages induced by radiotherapy, double strand breaks (DSB) are the most dramatic for the cells and their progeny. Eukaryotic cells have developed two main pathways of DSB repair with distinct mechanisms: non-homologous end-joining (NHEJ) and homologous recombination (HR). The NHEJ pathway is non-conservative and can operate DNA repair throughout the cell cycle, although it is most prevalent during G1/G0 phase [1, 2]. The NHEJ pathway proceeds by directly ligating the two ends of the break, which are protected by the binding of Ku80-Ku70 complex. HR is a highly conservative mechanism occurring mostly during S/G2 phase [2, 3] and relies on a homologous sequence to replicate the region lost during the DNA breakage event [4]. Defects in DNA repair pathways have been shown to be associated with strong sensitivity to ionizing radiations as observed in ataxia telangiectasia or Nijmegen breakage syndrome [5, 6]. A fraction of the general population (5 to 10%) is known to develop severe side effects after irradiation, suggesting heightened individual radiosensitivity [7]. The study of skin fibroblasts from such radiosensitive patients included in the COPERNIC cohort, has revealed several abnormal cellular features including impaired ATM nucleo-shuttling [8], lower clonogenic potential and delayed DNA repair leading to persistent DNA damages [9].

The mechanisms underlying these DNA repair impairments remain poorly understood. This study aimed to clarify the role of the transcription factor NFATc2 (Nuclear Factor of Activated T-cell), a protein we previously identified as severely downregulated in skin fibroblasts from radiosensitive patients, in the DNA repair defects observed in these cells [9]. Indeed, we also showed that NFATc2 protein deficiency observed in radiosensitive patients causes an accumulation of unrepaired DNA double-strand breaks 24h after 2Gy irradiation [9]. Interestingly Gabriel et al. reported by a large-scale proteomic approach that NFATc2 could interact with proteins involved in multiple DNA repair mechanisms [10]. Based on these results, we postulate that NFATc2 might play a role in DNA repair mechanisms through an as yet undetermined molecular process.

NFATc2 is one of the five members that constitute the NFAT protein family and is activated by the Ca^2+^/Calcineurin pathway. NFATc2 is a multifaceted protein and is involved in various biological processes such as inflammation, T cells activation [11] or cell cycle regulation [12]. Nevertheless, the involvement of NFATc2 in DNA repair has never been considered. In this study, we set out to characterize the interaction between NFATc2 and Ku80 involved in NHEJ repair pathway. We combined imaging and biochemical assays to demonstrate that NFATc2 binds Ku80 through its RHD domain. Finally, over-expression of NFATc2 in radiosensitive cells partially rescue their DNA repair defective phenotype. NFATc2 appears therefore as an important regulator of cellular radiosensitivity of skin fibroblasts.

## Materials & Methods

### Cell culture

Primary dermal fibroblasts were obtained from radiosensitive patients through the INSERM UMR1052 COPERNIC cell collection [13]. Patients were anonymized and provided informed consent in accordance with current ethical standards. This collection was approved by the regional Ethical Committee (CPP Sud-Est, Lyon, France) and cell lines were registered under the numbers DC2008-585 and DC2011-1437 with the Ministry of Research. The database derived from the COPERNIC collection is protected under the reference IDDN.FR.001.510017.000.D.P.2014.000.10300.

Primary normal human fibroblasts were obtained from skin surgical waste originating from Hospital Edouard Herriot in Lyon, France. Skin tissue samples were prepared and dispatched by DermoBiotec. Patients gave their informed consent. All healthy and patients’ fibroblasts were studied between P8 and P13. All treatments were performed on cells at confluency. The cell cultures were maintained as previously described [14].

### Cell irradiation

Primary Dermal Fibroblasts were irradiated with 2Gy using an XRAD320 X-Ray generator (Precision X-Ray, North Brandford, CT, USA) at a measured dose rate of 0.94 Gy/min, with a voltage of 320kV and current intensity of 12,5 mA.

### RNA interference, RNA extraction and Real-Time Quantitative PCR

Silencing of NFATc2 by lentiviral-mediated RNA interference was performed exactly as previously described [9]. mRNA extraction and RT-qPCR were performed as described [15]. Genes’ expression levels were normalized to *TBP* and *RPL13A* housekeeping genes expression levels using the 2^−ΔΔCt^ method. Primer list is available in supplementary data (Table S1)

### Immunofluorescence

Cells were cultivated and treated at confluency in µ-slide (#80807 Ibidi). Immunofluorescence were performed as previously described [9]. Primary antibodies (Table S2) were all incubated overnight at 4°C. After 3 washing steps with PBS, secondary antibodies (Table S2) were incubated for 1h at room temperature Images were acquired with a Nikon Tie motorized (Nikon). Images were treated and analyzed using ImageJ and QuPath softwares.

### Proximity ligation assay

Cells were cultivated and treated at confluency in µ-slide (#80807 Ibidi). Fixation and permeabilization were performed similarly as immunofluorescence assay. Primary rabbit anti-NFATc2, mouse anti-Ku80 antibodies, Rabbit anti-Ku70 and goat anti-53BP1 (Table S2), were incubated overnight at 4°C. The protocol for proximity ligation assay (#NC.MR.100 Red Navinci, Sweden) and co-staining was performed according to manufacturer recommendations. Images were acquired using a High Content screening microscopy CQ1 (Yokogawa, Tokyo, Japan), LSM980 Airyscan (ZEISS) and analyzed using ImageJ and QuPath softwares.

### Protein extraction and Western Blot analysis

Total proteins were extracted with RIPA buffer (R0278-500mL, Sigma-Aldrich), were quantified, loaded onto SDS polyacrylamide gels and transferred onto a PVDF membrane as previously described [14]. Membranes were blocked with a TBS 1X blocking solution containing 0.1% Tween20 and 5% BSA for 1h at RT. The membranes were incubated overnight at 4°C with the corresponding primary antibodies (Table S2). Membranes were incubated 1h at room temperature with HRP-Conjugated secondary antibodies (Table S2) and revealed with Super Signal West pico Plus (#34580 Thermo Fisher Scientific) using the Fusion FX system (Vilber). Signal quantification was achieved using Gel Analyzer (Version 23.1.1).

### Co-immunoprecipitation

HEK293T were co-transfected with vectors overexpressing FLAG-NFATc2, HA-Ku80 or with the controls vectors with JetPrime transfection reagent (Polyplus). Cells were irradiated at 2Gy and maintained at 37°C and 5% CO2 1h before lysis. Cells lysis and immunoprecipitation protocols are detailed in supplemental Materials and Methods. Antibodies references are available in supplementary data (Table S2).

### Truncated proteins production

The amino acid sequence of the RHD region of NFATc2 isoform c was determined using UniProt database (Q13469). The RHD region was identified within the NFATc2 transcript (*NM_173091*.*4)* using ApE software. The sequence coding for NFATc2 truncated for its RHD domain and the sequence coding for the RHD domain were both sent to VectorBuilder for plasmid production. HEK293T cells at 50% confluency were co-transfected with 500ng of each vector. Cells were irradiated at 2Gy and cell lysis was achieved 1h after exposure before performing co-immunoprecipitation.

### Stable overexpression of NFATc2

Fibroblasts from radiosensitive patients were transduced with a lentiviral vector overexpressing NFATc2. Lentiviral particles were produced by AniRA vectorology platform at SFR BioSciences Gerland, Lyon [16]. The detailed protocol for cells transduction and selection is given in supplemental Materials and Methods.

### Statistical analysis

Results are all expressed as mean ± SD. Statistical significance was assessed using Student’s t-test or Two-Way ANOVA using GraphPad Prism software (version 10.2.2). Results were considered statiscally significant when *p< 0.05, ** p<0.01.

## Results

As reported by Gabriel et al., NFATc2 may interact with proteins involved from various DNA repair pathways, including the Ku80 protein involved in NHEJ repair pathway [10]. In this study, we decided to mainly focus on this pathway as it predominates in dermal fibroblasts which are most often quiescent cells in the skin.

### NFATc2 interacts with the Ku80 protein in response to ionizing radiations

To determine whether the protein NFATc2 interacts with Ku80, we first performed a Proximity Ligation Assay (PLA) in control and patients’ fibroblasts, irradiated or not with 2Gy, to assess the spatial proximity between the two proteins. Without irradiation, we observed no or very few signals in both control and patient fibroblasts suggesting that NFATc2 does not interact with Ku80 in the absence of radiation exposure (**Figure 1A**). Interestingly, control fibroblasts showed a strong signal 1h after 2Gy irradiation compared to 0 Gy, indicating a marked proximity between the proteins NFATc2 and Ku80. This proximity was also observed in patients’ fibroblasts but to a lesser extent (**Figure 1A**), consistent with the previously detected downregulation of NFATc2 [9].

**Figure 1.**
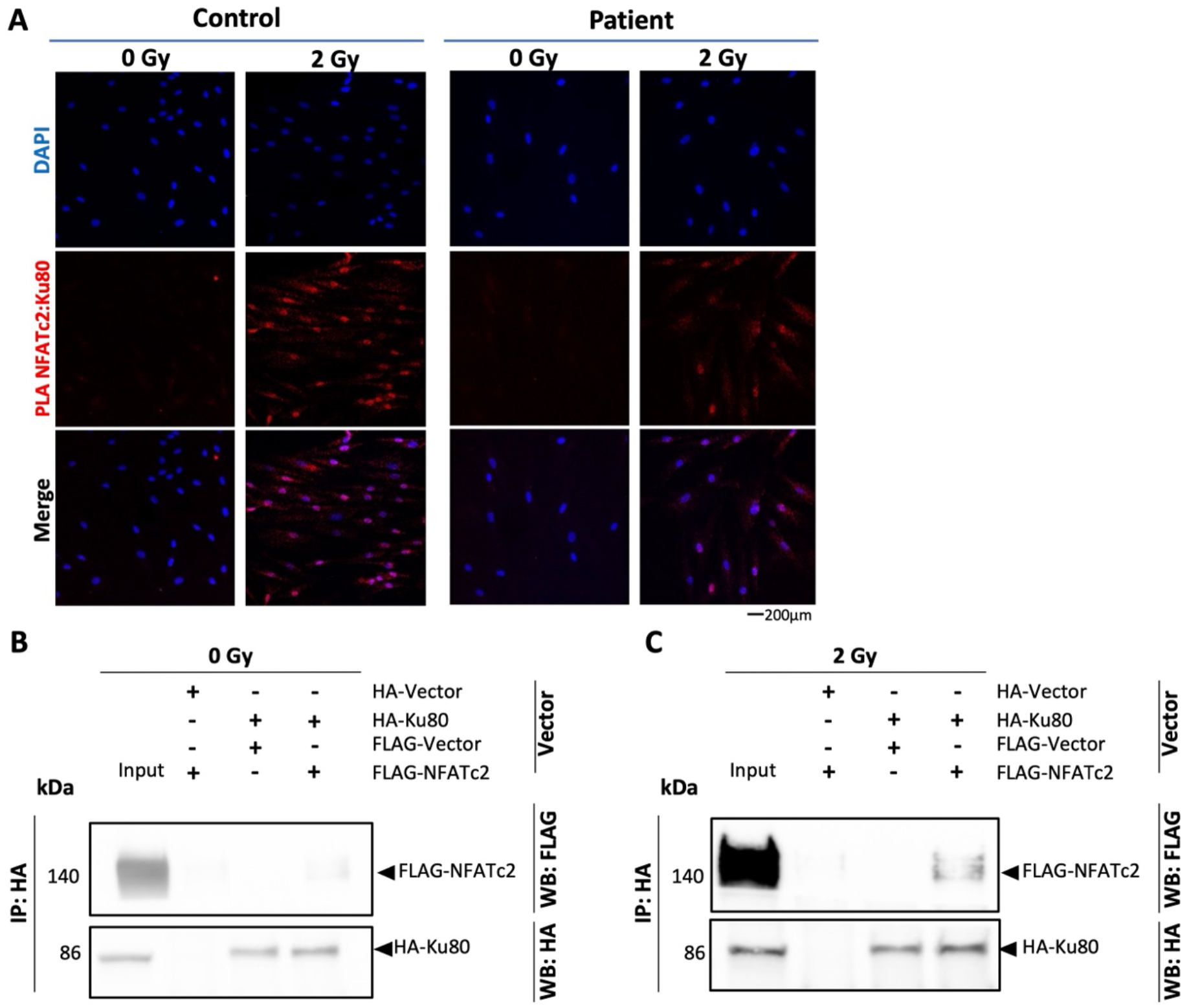
NFATc2 interacts with the protein Ku80. **(A)** Normal human primary fibroblasts (control) and fibroblasts from radiosensitive patients underwent a 2Gy irradiation. A Proximity Ligation Assay was performed 1h after exposure to ionizing radiations to test the interaction between NFATc2 and Ku80 (Red). HEK293T cells were co-transfected with a vector overexpressing HA tagged Ku80 (HA-Ku80) and FLAG tagged NFATc2 (FLAG-NFATc2). Empty HA (HA-Vector) and FLAG (FLAG-Vector) are used as negative controls. Co-Immunoprecipitation was performed on non-exposed cells **(B)** or 1h after a 2Gy irradiation **(C)**.

To further validate this interaction, we co-transfected HEK293T cells with vectors overexpressing FLAG-NFATc2 and HA-Ku80 and performed a co-immunoprecipitation assay 1h after irradiation. Little co-immunoprecipitation of FLAG-NFATc2 with Ku80 was observed in the non-exposed condition (**Figure 1B)**, whereas2 Gy exposure clearly enhances it **(Figure 1C)**. Taken together, these results demonstrate that NFATc2 interacts, directly or indirectly, with Ku80 in response to ionizing radiation. However, the NFATc2 domain mediating this interaction remained to be identified.

### NFATc2 RHD domain is required for the interaction with Ku80

To identify the NFATc2 domain involved in this interaction, we co-transfected HEK293T cells with a vector overexpressing HA-Ku80 together with vectors overexpressing either FLAG-NFATc2 lacking its Rel Homology Domain (RHD) (**Figure 2A**) or the FLAG-RHD alone. Here we show that Ku80 fails to co-immunoprecipitate with NFATc2 lacking its RHD domain (**Figure 2B**), but co-immunoprecipitated with RHD domain alone (**Figure 2C**), indicating that NFATc2 RHD region is required for the interaction with Ku80. However, the process by which NFATc2 regulates DNA repair, and particularly the NHEJ pathway, in response to radiation-induced DNA damage remains unclear at this stage.

**Figure 2.**
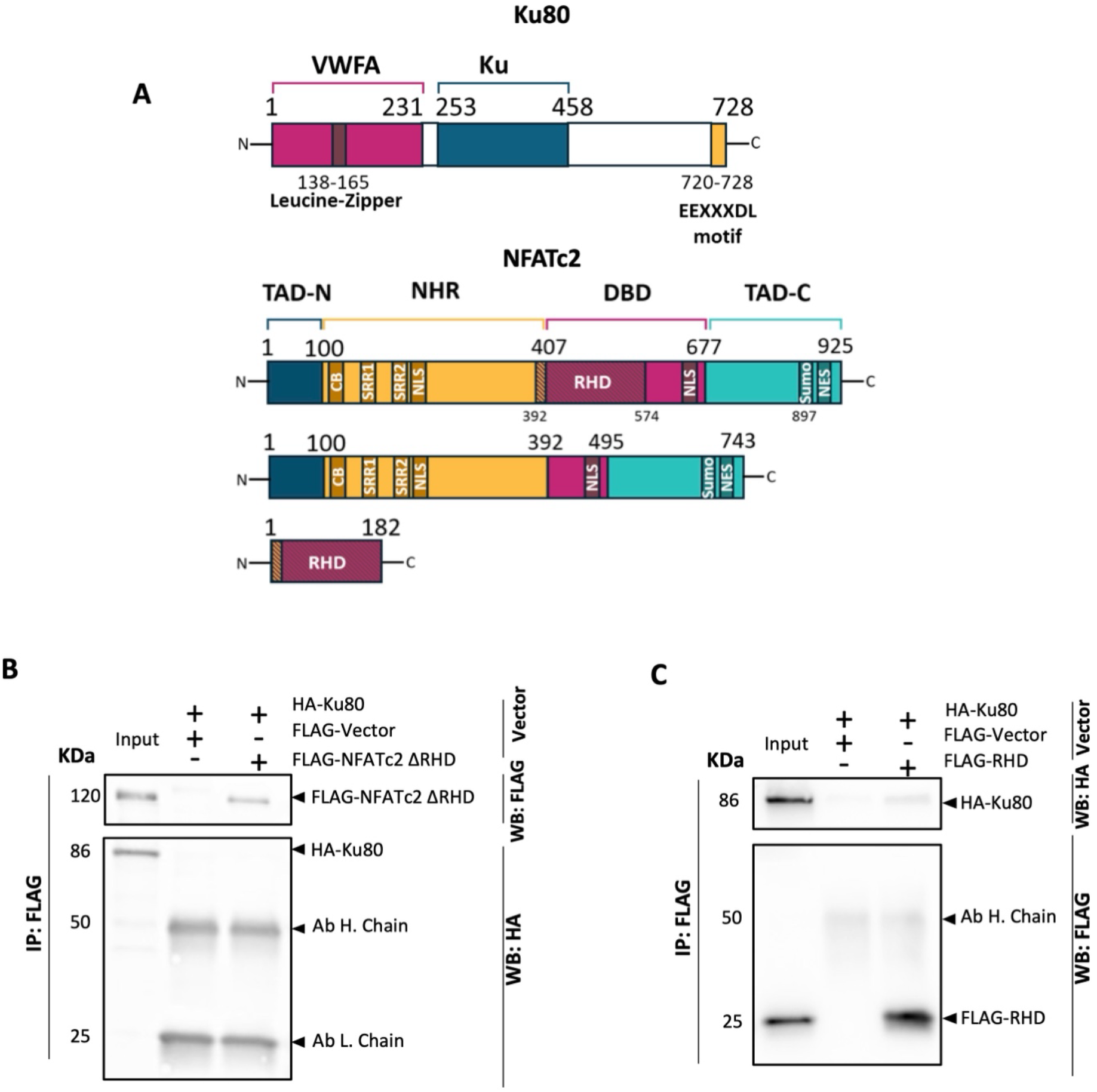
NFATc2 interacts with the protein Ku80 through its RHD domain. **(A)** Schematic representation of Ku80 domains and NFATc2 full-length domains (Top), NFATc2 truncated for RHD (Middle) and the RHD domain alone (Bottom). **(B)** HEK293T cells were co-transfected with vectors overexpressing the HA tagged Ku80 (HA-Ku80) and FLAG tagged NFATc2 protein truncated for its RHD domain (C) or with the FLAG tagged RHD domain only (FLAG-RHD) **(C)** in order to perform a co-immunoprecipitation assay 1h after a 2Gy irradiation. TAD: Transactivation Domain, NHR: NFAT Homology Region, DBD: DNA Binding Domain, CB: Calcineurin Binding, NLS: Nuclear Localization Signal, NES: Nuclear Export Signal, SRR: Serine Rich Regions

### NFATc2 NFATc2 and Ku80 complex does not colocalize on DNA double strand breaks

To better understand the role of NFATc2 in the regulation of NHEJ pathway, we first assessed the subcellular localization of the NFATc2:Ku80 complex over time in response to ionizing radiation. We combined PLA labelling of NFATc2:Ku80 with immunostaining of DNA double strand breaks using 53BP1, in normal primary fibroblasts irradiated at 2Gy and fixed at the indicated time post-irradiation (**Figure 3**). Interestingly, the interaction peaks between 30min and 1h after irradiation, then begins to decrease by 2h, even though double-strand breaks are still observable. Notably, NFATc2:Ku80 complex did not consistently colocalize with 53BP1 foci, suggesting that it may not be directly involved in NHEJ repair mechanism but rather in upstream DNA damage signaling.

**Figure 3.**
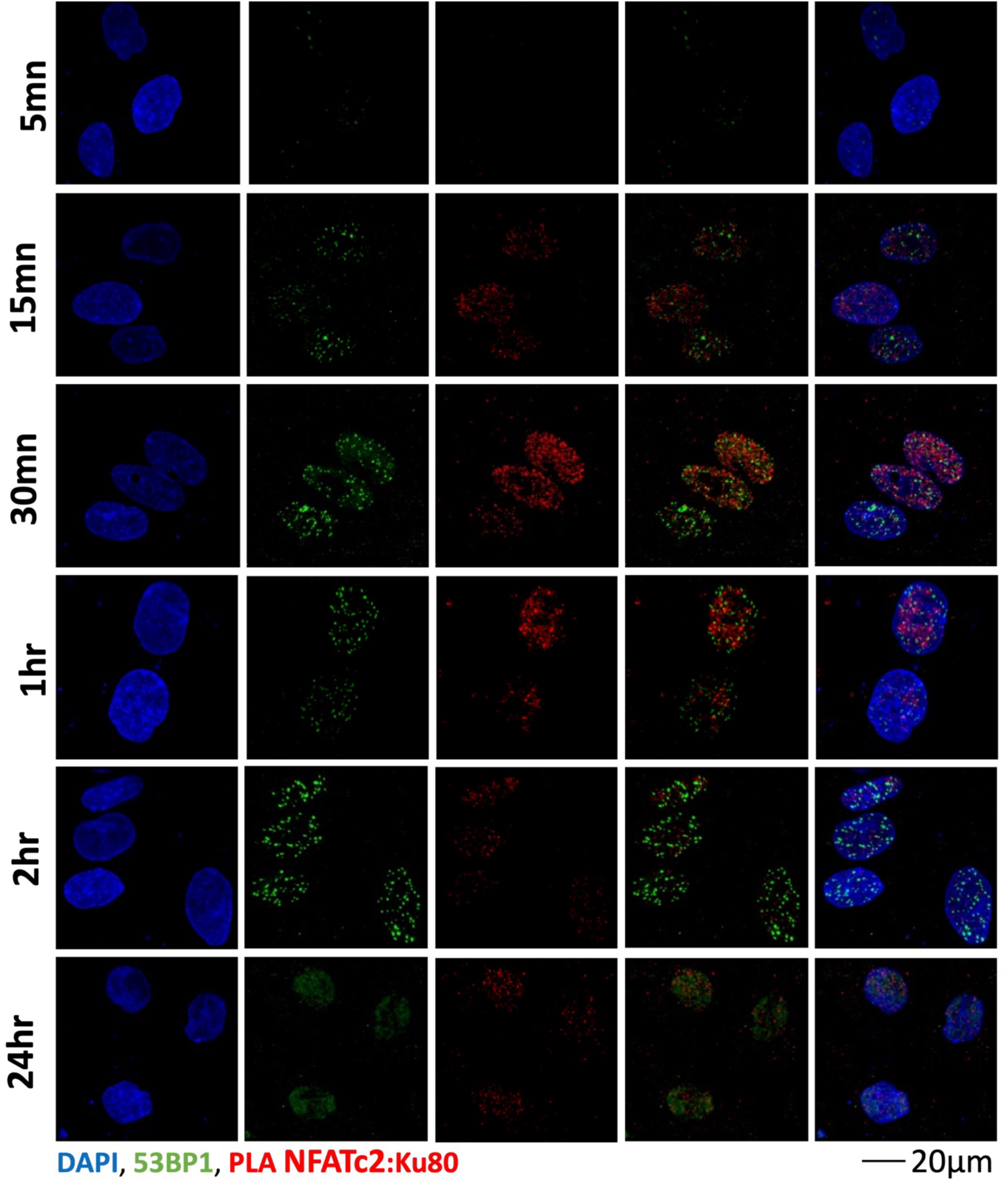
NFATc2 and Ku80 complex do not colocalize with DNA double-strand breaks. Normal fibroblasts were irradiated at 2Gy and fixed at indicated time after exposure. A Proximity Ligation Assay was performed against NFATc2 and Ku80 (Red) and co-immuno staining of 53BP1 was carried out to label DNA double-strand breaks (Green).

### NFATc2 is necessary to initiate the NHEJ pathway in response to ionizing radiations

To investigate the mechanism by which NFATc2 regulates DNA double-strand break repair, we analyzed the integrity of the NHEJ pathway by immunoblotting, 1h after 2Gy irradiation, in control fibroblasts and in fibroblasts silenced for *NFATc2* by RNA interference. We first confirm silencing at the RNA level by RT-qPCR and observe 70% decrease of *NFATC2* transcript in cells stably expressing the *NFATC2* shRNA (**Supplementary Figure 1**). Meanwhile, DNA-PKcs protein levels did not change significantly in response to ionizing radiation **(Figure 4A and B)**. However, DNA-PKcs phosphorylation was significantly induced in both control and NFATc2-silenced fibroblasts upon exposure, indicating that the NHEJ pathway is activated in response to ionizing radiation **(Figure 4C and D)**. This induction is approximately twofold lower in NFATc2-silenced fibroblasts **(Figure 4D)** thereby pointing to impaired NHEJ activation upon NFATc2 silencing in normal fibroblasts.

**Figure 4.**
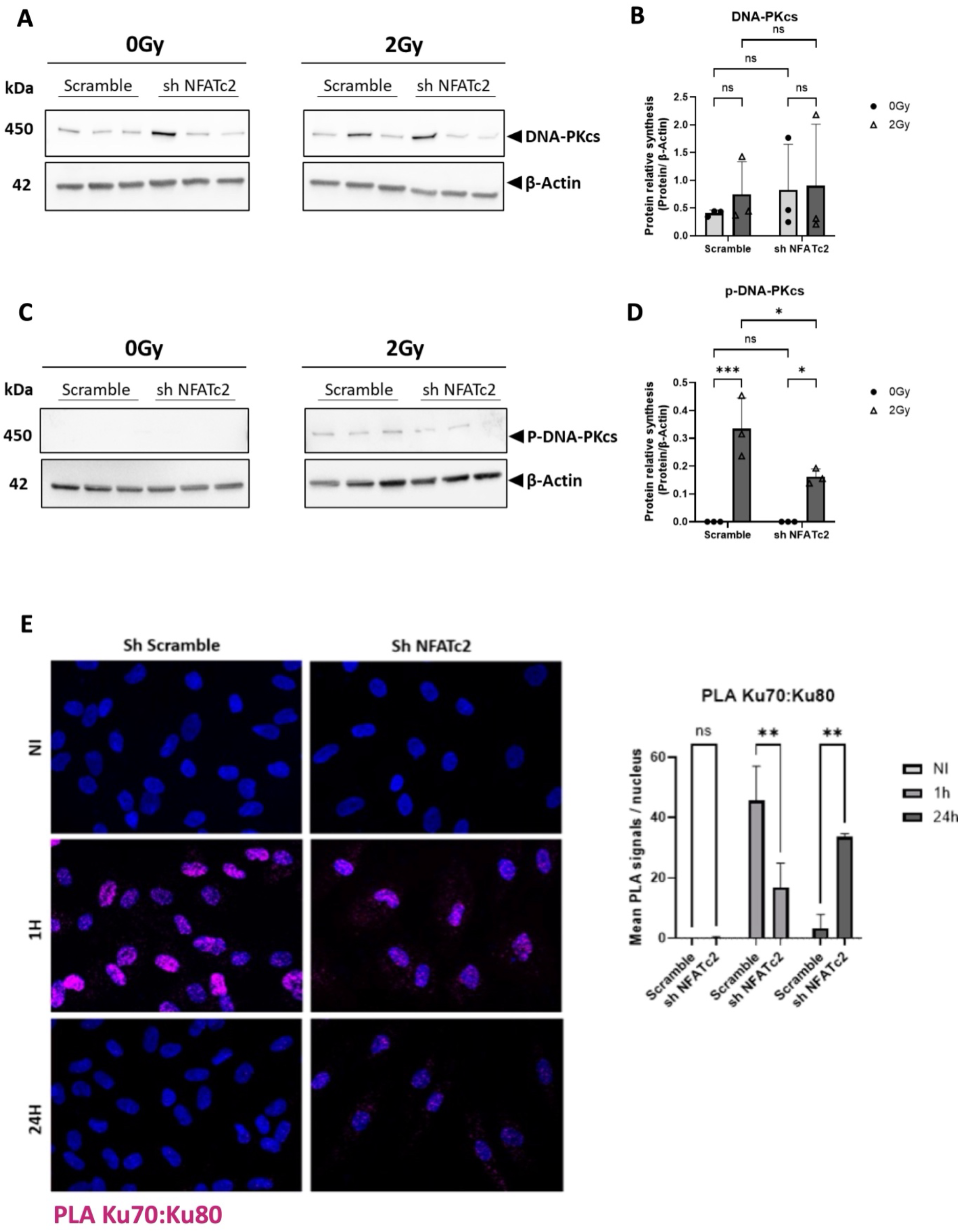
NFATc2 is necessary to induce DNA-PKcs phosphorylation and to allow normal Ku70/Ku80 interaction. Three different human normal fibroblasts strains were transduced with a control shRNA (Scramble) or NFATc2 targeting shRNA (sh NFATC2). The cells were irradiated at 2Gy and lysed 1h after exposure. Western Blot analysis was performed on non-exposed and irradiated cells to asses protein levels for DNA-PKcs **(A)** or P-DNA-PKcs **(C). (B)** Western Blot quantification for DNA-PKcs and **(D)** P-DNA-PKcs. Three independent experiments were performed (n = 3) for each condition. Results are mean ± SD. ANOVA2 with Tukey’s multiple comparison. *P <0.05; ns: P> 0.05. **(E)** Primary human fibroblasts transduced with a control or sh NFTAC2 were irradiated (IR) with 2 Gy or left untreated (NI) and then fixed 1 hour or 24 hours after irradiation to perform a PLA assay on the Ku70 and Ku80 proteins. The PLA signal is magenta. Quantification of PLA signals in cell nuclei is shown on the right panel. N=2. A minimum of 150 cells were analyzed per condition. ANOVA2 with Tukey’s multiple comparison ns: P> 0.05; ** P <0.01.

We next assessed the effect of NFATc2 silencing on the Ku70-Ku80 dimerization, an early step of NHEJ process. As expected, one hour after exposure, control fibroblasts show significantly more PLA signals than fibroblasts transduced with the NFATc2 shRNA (**Figure 4E)** indicating greater Ku70-Ku80 dimerization in the presence of NFATc2. Conversely, 24h after irradiation, very few PLA signals are detected in the scramble control, consistent with reduced NHEJ activity, whereas, in contrast, numerous PLA signals persist in fibroblasts transduced with the NFATc2 shRNA **(Figure 4E)**. Together, these results suggest a delay in the onset of NHEJ repair and/or slower repair in the absence of NFATc2, possibly due to reduced Ku70-Ku80 dimerization.

### NFATc2 over-expression in fibroblasts from radiosensitive patients restores DSB repair

Since NFATc2 is poorly expressed in radiosensitive patients’ cells, we examined how its re-expression affects DSB repair after X-irradiation. To do so, primary fibroblasts from five radiosensitive patients of the COPERNIC collection [13] (COP009, COP053, COP057, COP011 and COP049) were transduced with a lentiviral NFATc2 expression vector. A strong induction of the NFATc2 transcript was observed in the five cells strains transfected with the NFATc2 cDNA construction compared to empty vector **(Figure 5A)**, and NFATc2 overexpression was also confirmed at the protein level (**Figure 5B and 5C**). We then assessed the effect of this overexpression on the number of residual unrepaired DSBs 24h after irradiation, quantified by γH2AX foci counting after immunofluorescence staining. The number of residual foci increases in the 5 patients cell strains 24h after irradiation, particularly for COP053, COP011 and COP49 **(Figure 5D to 5H)**, that appeared to be the most radiosensitive based on this parameter. Upon NFATc2 overexpression (rescue condition), this increase of residual foci was abolished in COP53 **(Figure 5E)**, strongly reduced in COP011 and COP049 **(Figure 5G and 5H)**, limited in COP057 **(Figure 5F)** whereas no effect was observed in COP009 **(Figure 5D)**. Collectively, these data suggest that NFATc2 re-expression in NFATc2-low radiosensitive fibroblasts, can rescue the defect of DSB repair in the most radiosensitive patient cells.

**Figure 5.**
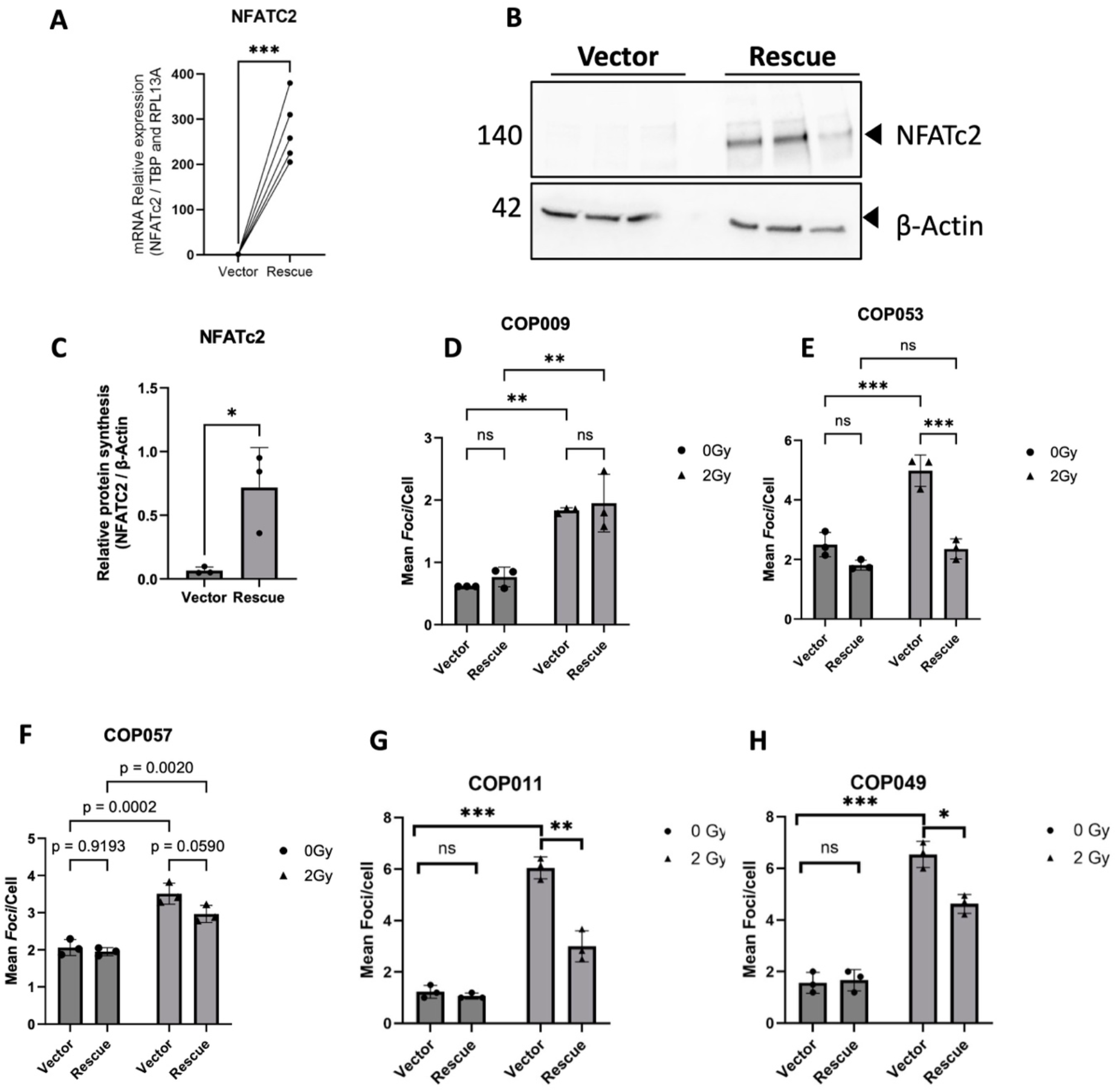
NFATc2 rescue in patients’ radiosensitive fibroblasts partially restores DNA double strand break repair activity. Fibroblasts from radiosensitive patients were transduced with a lentiviral vector overexpressing NFATc2. **(A)** RT-qPCR was performed to assess NFATc2 rescue, **(B)** Western Blotting analysis and **(C)** NFATc2 protein quantification in COP009, COP053, COP057. Rescued fibroblasts from 5 radiosensitive patients (COP009, COP053, COP057, COP011 and COP049) were irradiated at 2Gy and fixed 24h after exposure. P-γH2AX foci were stained by immunofluorescence and counted on 3 different fields for each strain **(D, E, F, G, H)** n>70 cells in total per condition. ANOVA2 with Tukey’s multiple comparison ns: P> 0.05; *P <0.05; ** P <0.01; *** P <0.001

## Discussion

The aim of this study was to provide a more detailed description of the DNA repair mechanisms triggered by exposure of fibroblasts to ionizing radiation in both healthy and radiosensitive context. Our previous work revealed a new function for the transcription factor NFATc2 in the response to radiation-induced DNA damage, showing that its silencing in normal fibroblasts was sufficient to generate cellular radiosensitivity and an accumulation of unrepaired DNA double-strand break [9].

To gain insight into the potential role of NFATc2 in regulating DNA repair, particularly the NHEJ pathway, we investigated its interaction with putative partners involved in DNA repair. We found that NFATc2 interacts with Ku80 within one hour of exposure to 2Gy radiation. Although the Ca2+/Calcineurin pathway responsible for NFATc2 activation and nucleo-shuttling could be triggered by radiation [17, 18], little evidence supports a direct causal link between radiation exposure and NFATc2 activation. Indeed, despite some study showing that radiations induce inflammation and activation of NFATc2 within T cells [19]; our findings extend these observations by suggesting a direct role for NFATc2 in DNA repair. Interestingly, it has been reported that inhibitors of the Ca^2+^/Calcineurin pathway, have been associated with an increased risk of skin cancer, resulting from DNA repair defects following UVB exposure [20, 21]. Conversely, several other transcription factors have been found to be involved in DNA repair by interacting with NHEJ actors. FOXL2 for instance, binds and sequesters Ku proteins; Their release from deacetylated FOXL2 allows assembly of the Ku complex at DSB sites [22]. Likewise, the EMT-inducing transcriptional repressor ZEB1 promotes NHEJ by direct interactions with 53BP1 [23]. These examples highlight the pleiotropic functions of certain transcription factors, which, beyond regulating transcription by direct binding to DNA, can modulate DNA repair mechanisms, particularly NHEJ, by interacting with DNA repair protein. PARP1 is another striking example as it controls multiple DNA repair pathways and regulates NFATc2 subcellular localization [24]. Moreover, NFATc2 is itself, a known PARP1 target in certain pathological contexts [25].

The interaction between NFATc2 and Ku80 involved the RHD domain of NFATc2. This domain is known to mediate not only NFATc2 binding to DNA, but also protein-protein interactions, by allowing dimerization of NFAT family proteins or their interaction with other cofactors [12]. Moreover, because the RHD is highly conserved across the NFAT family members (over 70% homology), other members likewise contribute to DNA repair. Consistent with this, NFAT5 (TonEBP) has been shown to interact with numerous DNA damage response proteins through its N-terminal RHD-containing region, to promote DNA damage tolerance, and to participate in R-loop resolution [26-28]. In contrast, no clear evidence links the other NFAT members to DNA repair mechanisms. Despite compelling evidence indicating the involvement of this domain in the interaction, further work is required to elucidate the exact interaction surface between NFATc2 and Ku80 and to identify the specific domain by which Ku80 interact with NFATc2. Another NFATc2 domain worth investigating would is the TAD-C domain which contains two Lysine residues, the Lys^684^ and the Lys^897,^ that can undergo SUMOylation [29]. This modification is specific to NFATc2 and appears to control NFATc2 its nuclear retention [29]. Notably, SUMOylation finely tunes multiple DNA repair mechanisms including NHEJ where it mediates DNA-PKcs activation and autophosphorylation [30].

Finally, by performing a functional rescue of NFATc2 in fibroblasts from radiation-sensitive patients with DNA repair defects, we obtained promising but heterogeneous results. We used lentiviral vectors to stably reintroduce expression of NFATc2. Our results indicates that the rescue was observed in 3 strains (COP053, COP011 and COP049), inconclusive in COP057 and with no discernible effect in COP009. It appears that the capacity for DNA repair is not uniform following exposure to ionizing radiation. The COP009 strain exhibited a significantly lower accumulation of additional breaks than the other 4 strains, with a 24-hour post-irradiation count of less than one additional DNA double strand-break. In contrast, the other strains exhibit a 2-to-3-fold change in the number of additional breaks following irradiation with up to 6 DNA double strand breaks. It is crucial to consider the number of accumulated and unrepaired breaks, which are the primary source of high genomic instability. Furthermore, it has been shown that unrepaired DNA damage can result in the establishment of an inflammatory niche and the emergence of a CAF-like phenotype in fibroblasts [31]. These results demonstrated the significant heterogeneity and difficulty in studying individual radiosensitivity. NFATc2 alone is probably not sufficient to fully restore repair capacity; rather, this also depends on the individual’s genetic background.

Our work reveals that NFATc2 down-regulation is not only a marker of radiosensitive patients’ fibroblasts but also a potential driver of radiation deleterious effects in these cells. By its interaction with Ku80, NFATc2 plays a key role in the initiation of the DNA repair, especially through the NHEJ pathway. The next step will be to study if this role of NFATc2 in DNA repair is limited to fibroblasts or can be found in other cell types and tissues. This would make NFATc2 a major regulator of the response to genotoxic stress, and thus a target of future treatments to restrain secondary effects of radiotherapy.

## Acknowledgements

We would like to warmly thank Dr Nicolas FORAY (INSERM U1296, Lyon-France) for providing us cell samples from the COPERNIC collection of radiosensitive patients.

We acknowledge the contribution of SFR Biosciences (Université Claude Bernard Lyon 1, CNRS UAR3444, Inserm US8, ENS de Lyon) [LyMic] especially Jacques BROCARD and Elodie CHATRE - for assistance with images acquisitions. We also thank Fabrice MURE for providing assistance and the formations necessary for cell irradiations using the XRAD-320.

We acknowledge the contributions of the CELPHEDIA Infrastructure (http://www.celphedia.eu/), especially the center AniRA and Caroline COSTA in Lyon for lentiviral particles production.

We acknowledge the contribution of PrImaTiss (Laboratoire de Biologie Tissulaire et d’Ingénierie Thérapeutique, UMR5305) especially Sandra FERRARO for her help with images acquisition.

---

**Supplemental Figure 1.**
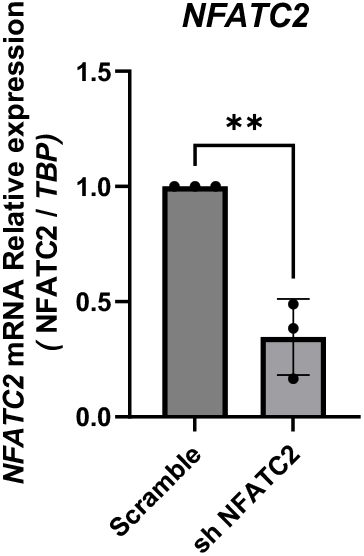

## References & citations

[1] Chiruvella KK, Liang Z, Wilson TE. Repair of double-strand breaks by end joining. Cold Spring Harb Perspect Biol. 2013;5:a012757.

[2] Mao Z, Bozzella M, Seluanov A, Gorbunova V. DNA repair by nonhomologous end joining and homologous recombination during cell cycle in human cells. Cell Cycle. 2008;7:2902–6.

[3] Karanam K, Kafri R, Loewer A, Lahav G. Quantitative live cell imaging reveals a gradual shift between DNA repair mechanisms and a maximal use of HR in mid S phase. Mol Cell. 2012;47:320–9.

[4] Scully R, Panday A, Elango R, Willis NA. DNA double-strand break repair-pathway choice in somatic mammalian cells. Nat Rev Mol Cell Biol. 2019;20:698–714.

[5] Pollard JM, Gatti RA. Clinical radiation sensitivity with DNA repair disorders: an overview. Int J Radiat Oncol Biol Phys. 2009;74:1323–31.

[6] Glover D, Harmer V. Radiotherapy-induced skin reactions: assessment and management. Br J Nurs. 2014;23:S28, S30-5.

[7] Lapierre A, Bourillon L, Larroque M, Gouveia T, Bourgier C, Ozsahin M, et al. Improving Patients’ Life Quality after Radiotherapy Treatment by Predicting Late Toxicities. Cancers (Basel). 2022;14.

[8] Sonzogni L, Granzotto A, Le Reun E, Al-Choboq J, Bourguignon M, Foray N, et al. Prediction of radiotherapy toxicity: 20 years of COPERNIC radiosensitivity diagnosis procedure. Cancer Radiother. 2024;28:435–41.

[9] Dulong J, Kouakou C, Mesloub Y, Rorteau J, Moratille S, Chevalier FP, et al. NFATC2 Modulates Radiation Sensitivity in Dermal Fibroblasts From Patients With Severe Side Effects of Radiotherapy. Front Oncol. 2020;10:589168.

[10] Gabriel CH, Gross F, Karl M, Stephanowitz H, Hennig AF, Weber M, et al. Identification of Novel Nuclear Factor of Activated T Cell (NFAT)-associated Proteins in T Cells. J Biol Chem. 2016;291:24172–87.

[11] Macian F. NFAT proteins: key regulators of T-cell development and function. Nat Rev Immunol. 2005;5:472–84.

[12] Mognol GP, Carneiro FR, Robbs BK, Faget DV, Viola JP. Cell cycle and apoptosis regulation by NFAT transcription factors: new roles for an old player. Cell Death Dis. 2016;7:e2199.

[13] investigators Cp, Granzotto A, Benadjaoud MA, Vogin G, Devic C, Ferlazzo ML, et al. Influence of Nucleoshuttling of the ATM Protein in the Healthy Tissues Response to Radiation Therapy: Toward a Molecular Classification of Human Radiosensitivity. Int J Radiat Oncol Biol Phys. 2016;94:450–60.

[14] Rorteau J, Chevalier FP, Bonnet S, Barthelemy T, Lopez-Gaydon A, Martin LS, et al. Maintenance of Chronological Aging Features in Culture of Normal Human Dermal Fibroblasts from Old Donors. Cells. 2022;11.

[15] Gonzalez Torres A, Chevalier FP, Aquino R, Aimard M, Baril P, Lamartine J. Spatiotemporal fluorescence imaging of microRNA activity in 3-D models of human epidermis reveals contribution of the Notch pathway in the regulation of miR-30a in aging skin. JID Innov. 2026;6:100444.

[16] Fusil F, Calattini S, Amirache F, Mancip J, Costa C, Robbins JB, et al. A Lentiviral Vector Allowing Physiologically Regulated Membrane-anchored and Secreted Antibody Expression Depending on B-cell Maturation Status. Mol Ther. 2015;23:1734–47.

[17] Yan J, Khanna KK, Lavin MF. Defective radiation signal transduction in ataxia-telangiectasia cells. Int J Radiat Biol. 2000;76:1025–35.

[18] Todd DG, Mikkelsen RB. Ionizing radiation induces a transient increase in cytosolic free [Ca2+] in human epithelial tumor cells. Cancer Res. 1994;54:5224–30.

[19] Tandl D, Sponagel T, Alansary D, Fuck S, Smit T, Hehlgans S, et al. X-ray irradiation triggers immune response in human T-lymphocytes via store-operated Ca2+ entry and NFAT activation. J Gen Physiol. 2022;154.

[20] Ume AC, Pugh JM, Kemp MG, Williams CR. Calcineurin inhibitor (CNI)-associated skin cancers: New insights on exploring mechanisms by which CNIs downregulate DNA repair machinery. Photodermatol Photoimmunol Photomed. 2020;36:433–40.

[21] Yarosh DB, Pena AV, Nay SL, Canning MT, Brown DA. Calcineurin inhibitors decrease DNA repair and apoptosis in human keratinocytes following ultraviolet B irradiation. J Invest Dermatol. 2005;125:1020–5.

[22] Jin H, Lee B, Luo Y, Choi Y, Choi EH, Jin H, et al. FOXL2 directs DNA double-strand break repair pathways by differentially interacting with Ku. Nat Commun. 2020;11:2010.

[23] Genetta TL, Hurwitz JC, Clark EA, Herold BT, Khalil S, Abbas T, et al. ZEB1 promotes non-homologous end joining double-strand break repair. Nucleic Acids Res. 2023;51:9863–79.

[24] Valdor R, Schreiber V, Saenz L, Martinez T, Munoz-Suano A, Dominguez-Villar M, et al. Regulation of NFAT by poly(ADP-ribose) polymerase activity in T cells. Mol Immunol. 2008;45:1863–71.

[25] Mou K, Zhou Y, Mu X, Zhang J, Wang L, Ge R. PARP1 Is a Prognostic Marker and Targets NFATc2 to Promote Carcinogenesis in Melanoma. Genet Test Mol Biomarkers. 2022;26:503–11.

[26] Kang HJ, Park H, Yoo EJ, Lee JH, Choi SY, Lee-Kwon W, et al. TonEBP Regulates PCNA Polyubiquitination in Response to DNA Damage through Interaction with SHPRH and USP1. iScience. 2019;19:177–90.

[27] Kang HJ, Cheon NY, Park H, Jeong GW, Ye BJ, Yoo EJ, et al. TonEBP recognizes R-loops and initiates m6A RNA methylation for R-loop resolution. Nucleic Acids Res. 2021;49:269–84.

[28] Ye BJ, Kang HJ, Lee-Kwon W, Kwon HM, Choi SY. PARP1-mediated PARylation of TonEBP prevents R-loop-associated DNA damage. DNA Repair (Amst). 2021;104:103132.

[29] Terui Y, Saad N, Jia S, McKeon F, Yuan J. Dual role of sumoylation in the nuclear localization and transcriptional activation of NFAT1. J Biol Chem. 2004;279:28257–65.

[30] Bhachoo JS, Garvin AJ. SUMO and the DNA damage response. Biochem Soc Trans. 2024;52:773–92.

[31] Legrand AJ, Poletto M, Pankova D, Clementi E, Moore J, Castro-Giner F, et al. Persistent DNA strand breaks induce a CAF-like phenotype in normal fibroblasts. Oncotarget. 2018;9:13666–81.

