## Supplemental methods table figure for "NFATc2 potentiates DNA double-strand breaks repair by interaction with Ku80 in radiotherapy patients"

**Supplemental Materials, Figures and tables**

**Supplementary materials and methods**

**Co-immunoprécipitation**

After HEK293T cells transfection and irradiation, a gentle lysis was performed with a lysis buffer containing 50mM Tris HCL, 150mM NaCl, 2mM EDTA, 1% NP-40 for 1h on ice. Lysates were centrifuged at 13000 rpm for 15min at 4°C and total proteins were quantified using Pierce BCA protein assay (#23225, ThermoFisher). 500µg of proteins were used for immunoprecipitation experiments. 20 µL Protein G Magnetic Beads (#70024 Cell Signaling Technology) were washed 3 times with lysis buffer. Cell lysates were incubated with 20µL of beads with rotation for 20min at RT in order to pre-clear the samples. 20 µL of protein G magnetic beads were conjugated with 3µg of anti-FLAG or 5µg of anti-HA antibodies for 1h30 at 4°C. Western Blot analysis was performed as described above. In order to remove aspecific signal due to antibody light chains masking our signal of interest, some IP were performed using anti-FLAG or anti-HA antibodies and membranes were revealed using a conformation specific secondary antibody.

**Fibroblasts transduction with lentiviral vectors for functional rescue**

Cells were seeded at 5000 cells/cm² and infected the next day with the lentiviral particles using a MOI of 25 and incubated overnight at 37°C and 5% CO2 with DMEM (Gibco) supplemented with 10% decomplemented serum. 24h after infection, the culture media was discarded and replaced with fresh complete culture medium. Cells were cultivated for 48h before selection. The selection was achieved using puromycin at 0.7µg/mL for 4 days and the cells were amplified for 1 week before performing experiments.

**Supplementary figures and tables**

**Table S1:** Primers used for RT-qPCR experiments

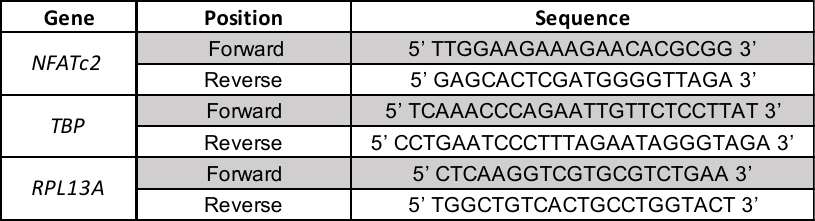

**Table S2:** Antibodies used and their respective application in the project

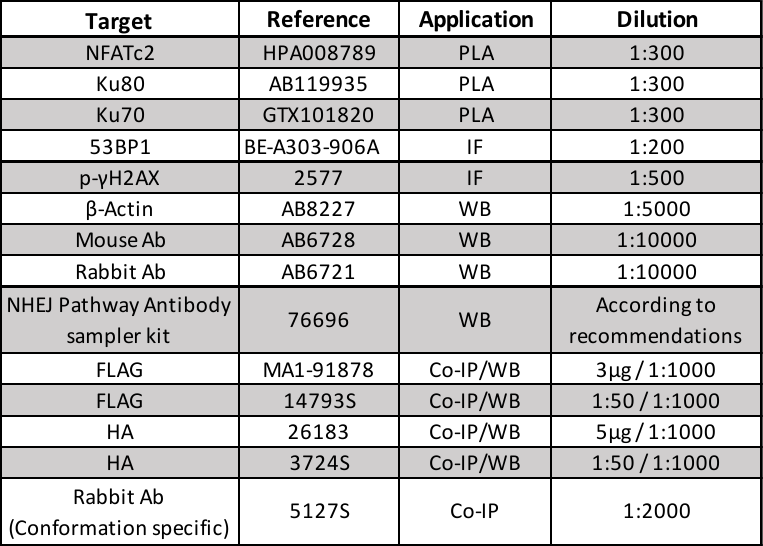

**Supplemental Figure 1.** Normal human fibroblasts (n=3) were transduced with a shRNA targeting NFATc2. RT-qPCR was performed to assess NFATc2 silencing. Results are mean ± SD. Unpaired Student t-test, ** p<0,01.

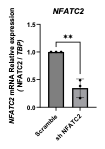
